# Geosmin suppresses defensive behaviour and elicits unusual neural responses in honey bees

**DOI:** 10.1101/2021.10.06.463314

**Authors:** Florencia Scarano, Mukilan Deivarajan Suresh, Ettore Tiraboschi, Amélie Cabirol, Morgane Nouvian, Thomas Nowotny, Albrecht Haase

**Author notes:** Correspondence to: Albrecht Haase. Department of Fundamental Microbiology, University of Lausanne, Lausanne, Switzerland (Amélie Cabirol).

## Abstract

Geosmin is an odorant produced by bacteria in moist soil. It has been found to be extraordinarily relevant to some insects, but the reasons for this are not yet fully understood. Here we report the first tests of the effect of geosmin on honey bees. A stinging assay showed that the defensive behaviour elicited by the bee’s alarm pheromone is strongly suppressed by geosmin. Surprisingly, the suppression is, however, only present at very low geosmin concentrations, and completely disappears at higher concentrations. We investigated the underlying mechanisms of the behavioural change at the level of the olfactory receptor neurons by means of electroantennography and at the level of the antennal lobe output via calcium imaging. Unusual effects were observed at both levels. The responses of the olfactory receptor neurons to mixtures of geosmin and the alarm pheromone component isoamyl acetate (IAA) were lower than to pure IAA, suggesting an interaction of both compounds at the olfactory receptor level. In the antennal lobe, the neuronal representation of geosmin showed a glomerular activation that decreased with increasing concentration, correlating well with the concentration dependence of the behaviour. Computational modelling of odour transduction and odour coding in the antennal lobe suggests that a broader than usual activation of different olfactory receptor types by geosmin in combination with lateral inhibition in the antennal lobe could lead to the observed non-monotonic increasing-decreasing responses to geosmin and thus underlie the specificity of the behavioural response to low geosmin concentrations.

## Introduction

Geosmin is a musty, earthly-smelling compound produced by multiple microorganisms from various clades such as cyanobacteria, actinobacteria (*e.g. Streptomyces sp*.), protozoa, moulds and fungi (Gerber and Lechevalier, 1965; Liato and Aïder, 2017). Actinobacteria (including *Streptomyces*) are widely associated with hymenopteran nests, which they likely protect from pathogens thanks to their natural production of antibiotics (Madden et al., 2013; Matarrita-Carranza et al., 2017; Rodríguez-Hernández et al., 2019). A recent study found that fire ant queens (*S. invicta*) preferentially started new nests in actinobacteria-rich soil. This attraction was mediated in part by geosmin and resulted in a higher survival rate of the queen (Huang et al., 2020). Geosmin is also ecologically important for other insects, such as the vinegar fly *Drosophila melanogaster* and the mosquito *Aedes aegypti*. However, it evokes dramatically different responses in those species. Geosmin elicits a strong aversion in *D. melanogaster*, even in the presence of attractive compounds (Becher et al., 2010; Stensmyr et al., 2012). This could be to avoid oviposition on mouldy, unsuitable fruit (Stensmyr et al., 2012), or to provide a better contrast with fallen ripe fruits, thus making the search more efficient (Galizia, 2020). In *Ae. aegypti*, on the contrary, geosmin is a strong attractant (Melo et al., 2020). This is likely because it signals the presence of wet soil, in which eggs can be laid. Indeed, geosmin is also known for being one of the main components of Petrichor, "the smell of wet soil" (Liato and Aïder, 2017).

In the first olfactory processing brain centre of insects, the antennal lobe (AL), most odours elicit a combinatorial pattern of glomerular responses (Joerges et al., 1997; Haverkamp et al., 2018). However, some odours, often with high biological relevance (e.g. sex pheromones), only activate a single glomerulus. When this is followed by segregated processing also in higher brain centres, resulting in stereotypical responses, this circuit is termed a labelled line (Haverkamp et al., 2018). Geosmin is one of very few compounds that activate only a single glomerulus in *D. melanogaster* and *Ae. Aegypti* mosquitoes (Stensmyr et al., 2012; Melo et al., 2020). In flies, its processing is further functionally segregated in higher brain centres, where it takes priority over other olfactory signals to trigger an avoidance behaviour (Stensmyr et al., 2012).

Despite the ecological relevance of geosmin in many insect species, behavioural and physiological responses to this odour have not been investigated in the honey bee *Apis mellifera* yet. Here, we provide the first data tackling this question. Within the behavioural repertoire of honey bees, we focus on its defensive response. Honey bees defend their colony by stinging potential intruders, a behaviour that is stimulated in the presence of the alarm pheromone released by other colony members (Nouvian et al., 2016). There is some evidence that honey bees become more defensive in high relative humidity in the field, such as is found after rain (Brandeburgo, M. M Goncalves and Kerr, 1982; Southwick and Moritz, 1987). Since geosmin is also released from the soil after rain, we wondered if it could be responsible for this modulation of the defensive response. We thus tested this hypothesis using a well-established stinging assay (Nouvian et al., 2015). Furthermore, we searched for neuronal correlates for the behaviour at the level of the antennal lobe projection neurons via *in vivo* two-photon calcium imaging (Haase et al., 2011) of geosmin-elicited activity, and at the level of the olfactory receptor neurons via electroantennography. Finally, we used a spiking neural network model to investigate how the observed partially unusual neuronal responses relate to the current understanding of the bee olfactory system.

## Results

### Behavioural responses to geosmin, IAA, and their mixtures

The stinging behaviour of a dyad of bees towards a yellow-black rotating dummy, presented inside an arena was monitored under the exposure to different odour stimuli. In a control group, bees were exposed to pure mineral oil. In this group, in 15% of the trials, at least one of the two bees showed stinging behaviour against the dummy (**Figure 1A, Video 1**). If bees were exposed to geosmin within the arena, no effect on the frequency of stinging behaviour was observable. If instead bees were exposed to isoamyl acetate (IAA, at a concentration of 10^−1^), an active compound of the bees’ alarm pheromone (Boch et al., 1962), the stinging behaviour increased significantly to 50% stinging (*t* (47) = 3.3, *p* = 0.002) as expected (**Figure 1B**).

**Figure 1.**
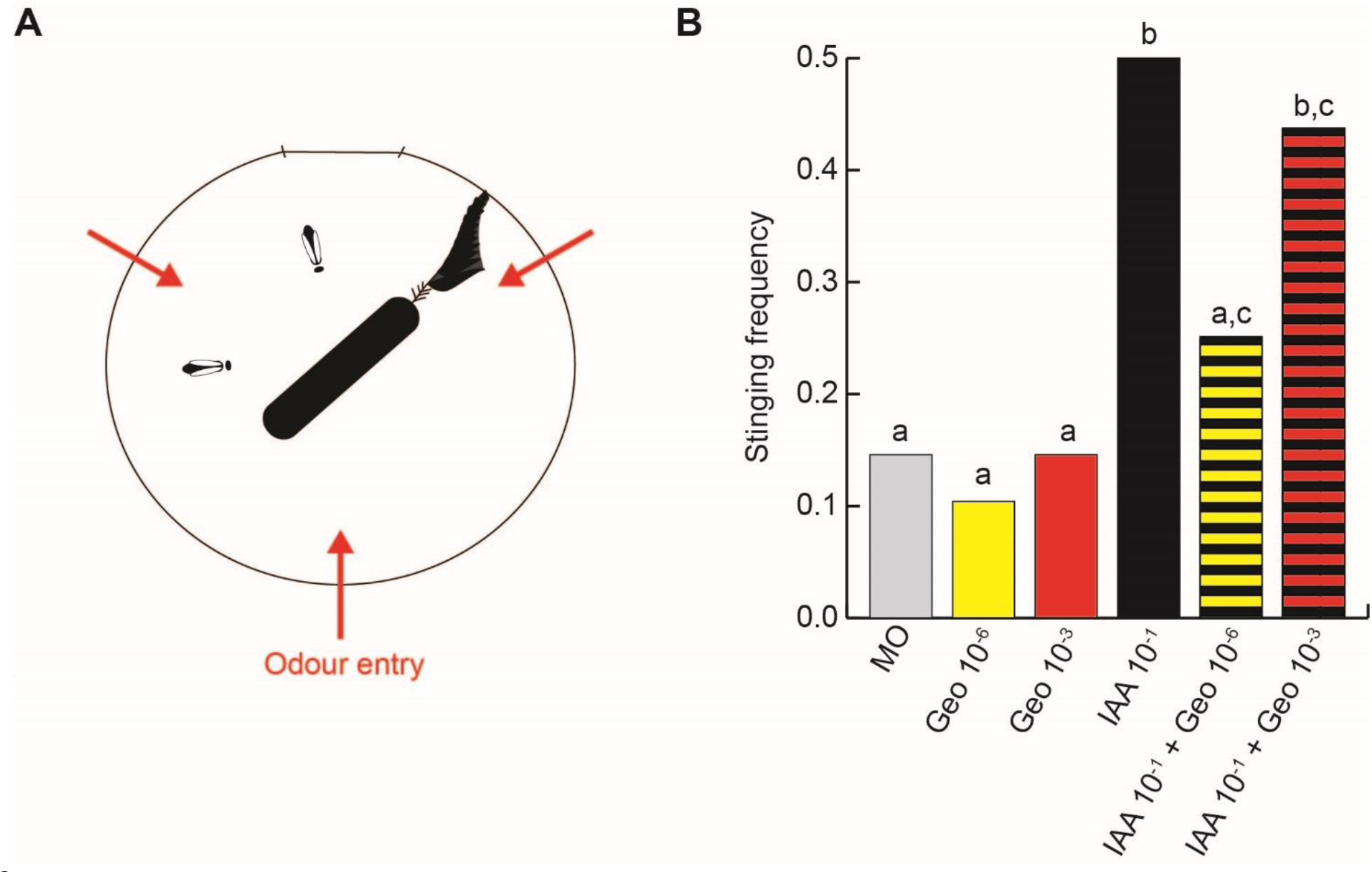
A low concentration of geosmin prevents recruitment into stinging behaviour by IAA. **A.** Schematic of the behavioural assay. A dyad of bees was presented with a rotating dummy, that they could choose to sting or not. The red arrows denote the entry points of the airflow carrying the odours into the arena. **B.** Frequency of trials in which at least one of the bees exhibited stinging behaviour (*n* = 48 dyads of bees per group). MO: mineral oil (solvent control); Geo 10^−x^: geosmin at concentration 10^−x^; IAA 10^−1^: isoamyl acetate at concentration 10^−1^. The groups labelled with the same letter are not significantly different from each other (GLM, corrected for multiple comparisons with an FDR procedure).

When bees were exposed to a mixture of IAA and a low concentration of geosmin (10^−6^), clear signs of an interaction between both stimuli were observable with a stinging probability of 25%, a significant reduction of the frequency of this behaviour compared with bees exposed to IAA only (*t* (47) = 2.4, *p* = 0.02). There was no significant difference anymore with respect to the mineral oil control (*t* (47) = 1.1, *p* = 0.27, **Figure 1B**).

However, when the geosmin concentration in the mixture was 10^−3^, stinging was observed in 40% of the cases, which was not significantly different from the behaviour of the bees stimulated with IAA only (*t* (47) = 0. 6, *p* = 0.57). Indeed, this stinging frequency was again significantly higher than for control bees (*t* (47) = 2.9, *p* = 0.006), suggesting that the behaviour was driven by the response to IAA in this case (**Figure 1B**).

### Olfactory receptor neuron responses at the level of the antennas

We started to search for the neuronal correlates of the behavioural effect of Geosmin combined with IAA at the periphery of the olfactory system, targeting the olfactory receptor neurons (ORNs) in the antennas. Electroantennography measures a voltage change between electrodes at each end of the antenna in response to an odour exposure. This signal has an amplitude proportional to the sum of the activity elicited in all ORNs. A typical signal in response to IAA is illustrated in **Figure 2A**. In our experiment, honey bee antennas (n=24) were exposed to Geosmin at 4 logarithmically varying concentrations from 10^−6^ to 10^−3^, to two concentrations of IAA: 10^−3^ and 10^−1^, and to mixtures of both odours at all these concentrations. Results showed a response significantly different from the mineral oil control at a geosmin concentration of 10^−5^ or higher. The signals followed the typical exponential amplitude increase in response to growing concentrations (**Figure 2B**). The response to the lowest concentration of geosmin (10^−6^) was not significantly different from the control, probably because the signal intensity was too little to be detected with this method.

**Figure 2.**
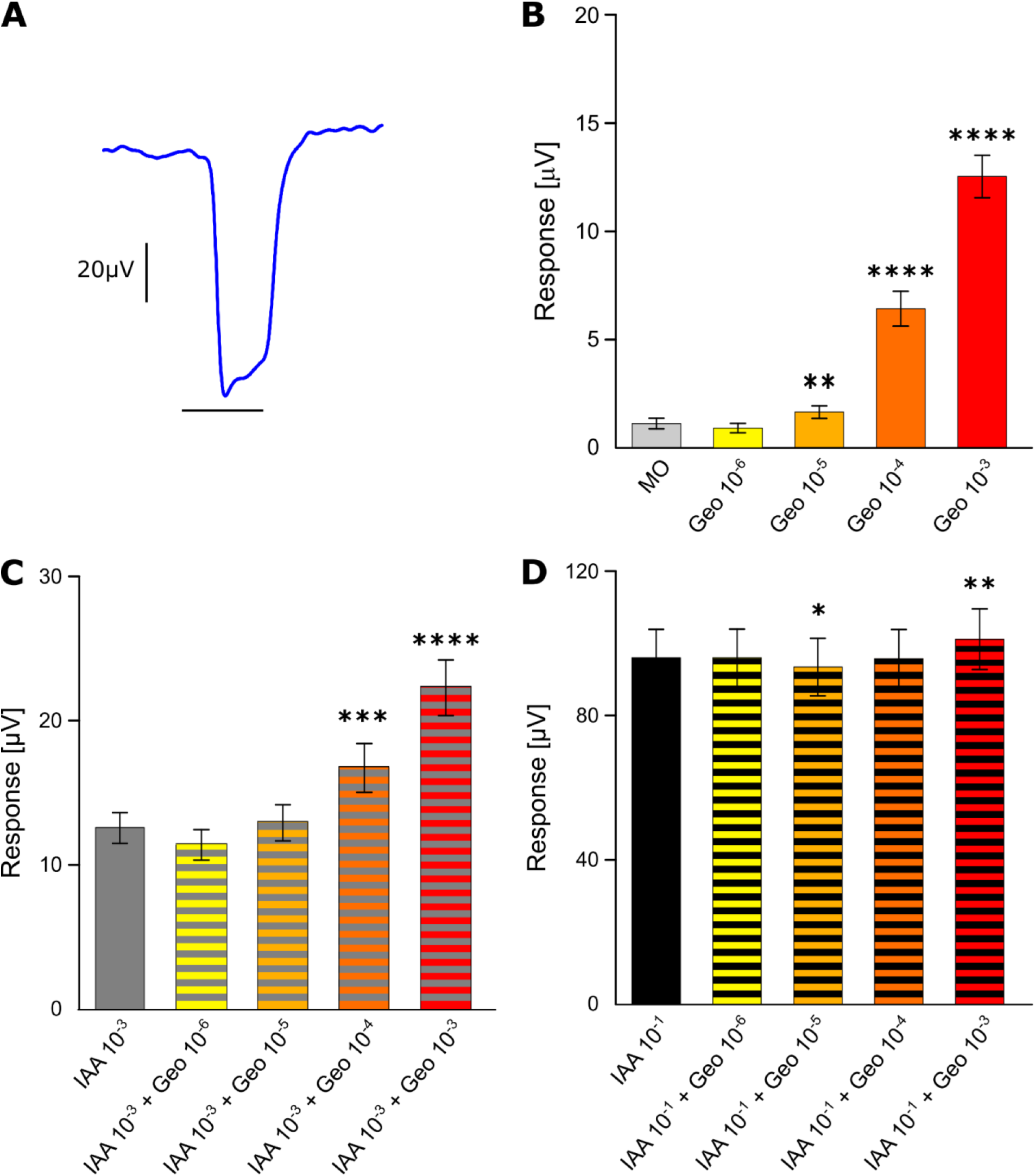
Olfactory receptor neuron responses. **A.** Example of a typical EAG response curve to IAA 10^−1^. The black line below the recording indicates the stimulus presence (2 s). **B.** Mean response amplitude (±SEM) to different concentrations of geosmin. **C.** Mean response amplitude (±SEM) to different concentrations of geosmin combined with IAA 10^−3^. **D.** Mean response amplitude (±SEM) to different concentrations of geosmin combined with IAA 10^−1^. *IAA 10^−x^*: Isoamyl acetate 10^−x^ (vol/vol); *Geo 10^−x^*: Geosmin 10^−x^ (vol/vol); Statistics are Bonferroni corrected comparisons with Control, IAA3, and IAA1, in B, C, D respectively: ****: *p* < 0.0001; ***: *p* < 0.001; **: *p* < 0.01; *: *p* < 0.05.

Stimulation with IAA at a concentration of 10^−3^ induced a strong response, which seemed reduced in the presence of Geosmin 10^−6^ although the effect is not significant. With Geosmin at concentrations 10^−4^ or higher, the signals in response to mixtures were significantly increased in comparison to IAA-only stimulation (**Figure 2C**). When the antennas were exposed to mixtures with IAA at a concentration of 10^−1^, the signals showed a reduction when a Geosmin concentration of 10^−5^ was present although in this case, the difference was significant with respect to IAA (*p* = 0.037, **Figure 2D**).

### Projection neuron responses

To trace the odour-induced activation across the next processing level, we performed calcium imaging of the projection neurons in the honey bee antennal lobes. These neurons convey information from the ALs to higher-order brain centres. Imaging experiments were performed on 14 bees exposed to the same odours at the same concentrations as in the behavioural assay. The change in fluorescence induced by the odour stimulus was recorded in 19 glomeruli (**Figure S1**). When averaged over the 3 s stimulus period, these glomerular signals showed highly stereotypical response patterns across bees (**Figure 3C** shows the IAA geosmin mixture as an example).

**Figure 3.**
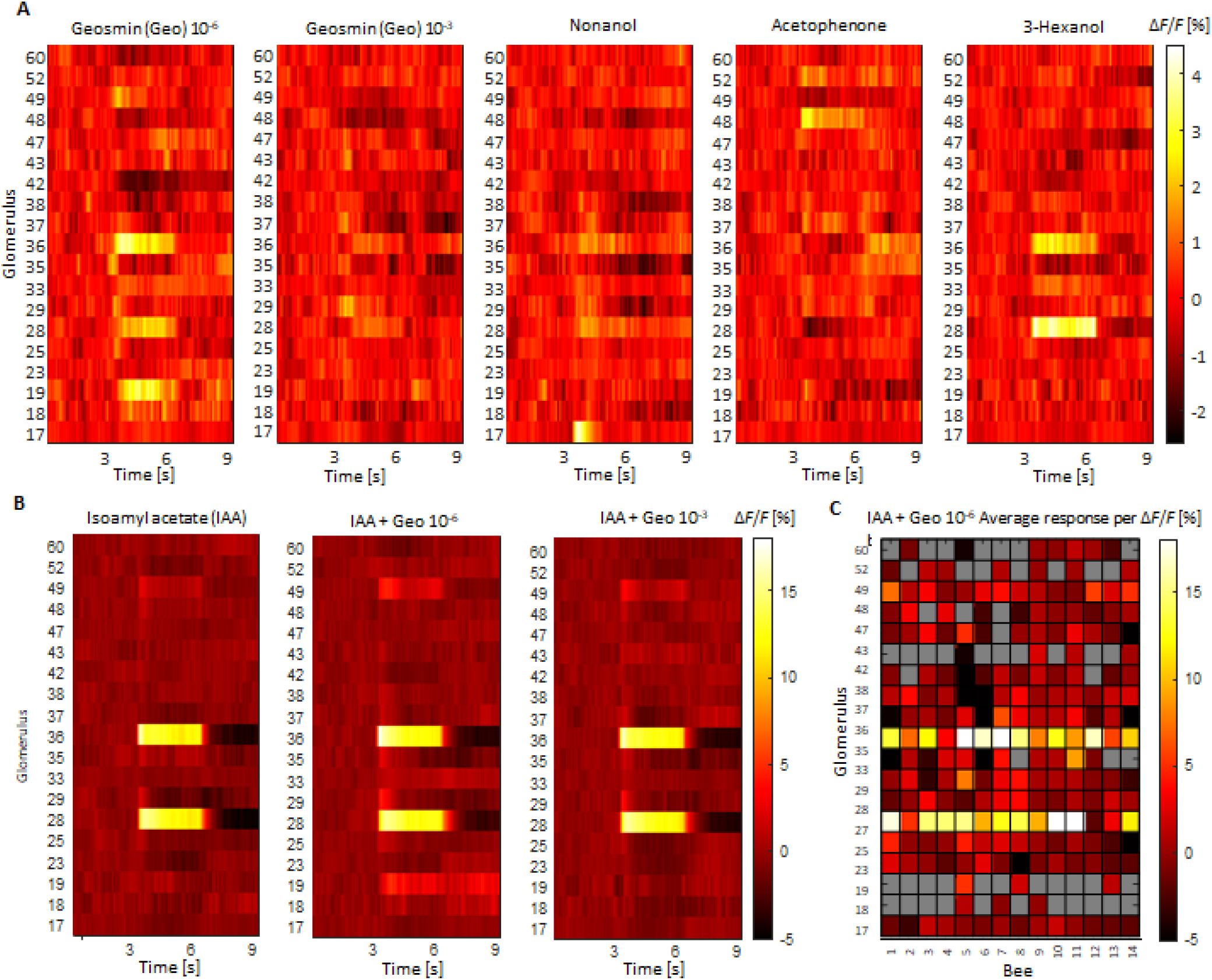
Glomerular responses. **A.** Mean glomerular response curves to pure Geosmin in two different concentrations, 10^−6^ and 10^−3^, and to three floral odour compounds; 1-nonanol, benzaldehyde, and 3-hexanol at concentration 5∙10^−3^ for 19 glomeruli (**Figure S1**) averaged over all identified glomeruli in 14 bees. Odour stimulation started after 3s and lasted 3s. Color-coded is the calcium-induced fluorescence change in percent. **B.** Mean glomerular response curves to the alarm pheromone compound isoamyl acetate (IAA) at 10^−1^ concentration and to mixtures of IAA with the 2 concentrations of geosmin. The colour-scale is different, given the stronger responses to 10^−1^ IAA. **C.** Time-averaged glomerular response amplitudes to the mixture of IAA and 10^−6^ geosmin in each bee, showing the stereotypy of the response. Grey represents cases in which glomeruli were not clearly identified.

The signal’s contrast is increased by averaging across subjects. The average temporal response curves show the highly different responses to 4 pure odours (**Figure 3A**): Geosmin at the same two concentrations as used in the behavioural tests, 10^−6^ and 10^−3^ in mineral oil, and the three floral odours 1-nonanol, benzaldehyde, and 3-hexanol all at a concentration of 5∙10^−3^ in mineral oil. Geosmin at concentration 10^−6^ elicits responses in various glomeruli, the spectrum might be slightly broader than those of the floral odour showing responses in 6 glomeruli, while the floral odours elicit responses in up to 4 glomeruli. Surprisingly, these geosmin responses disappear almost completely at the 100-fold higher concentration of 10^−3^, suggesting a non-monotonic concentration dependence in the PN response pattern.

Next, bees were stimulated with the alarm pheromone compound IAA again at the same concentration as in the behavioural experiment of 10^−1^, which is at least 20-fold higher than the other stimuli and accordingly elicits much stronger responses. In addition to IAA, also its mixtures with geosmin were tested. Responses seem to be accurately the sum of the single compound responses. Noticeably, all glomeruli sensitive to IAA, seem also to be sensitive to geosmin. Also in the mixtures, the geosmin contributions are clearly visible at a concentration of 10^−6^ but almost completely disappear at a concentration of 10^−3^.

### Computational modelling

The observation of PN responses in the AL that are non-monotonic as a function of concentration is unusual. We have built a computational spiking neural network model of the early olfactory system of bees in order to investigate whether this and our other observations are consistent with our current understanding of the system. The model builds on earlier works (Nowotny et al., 2013; Chan et al., 2018) and describes olfactory receptors with a two-stage binding and activation process (Rospars et al., 2008). Olfactory receptors then excite olfactory receptor neurons (ORN), which in turn excite projection neurons (PNs) and local neurons (LNs) in the antennal lobe. All ORNs with the same receptor type project to the same glomerulus (Vosshall et al., 2000). LNs inhibit PNs and LNs in all other glomeruli. The circuit is illustrated in **Figure 4A**. The model has 160 receptor types and hence glomeruli, 60 ORNs of each type, and 5 PNs and 25 LNs per glomerulus. Further details are given in the Methods. **Figure 4C** shows an example data trace from the simulations. Odours were introduced as a step change from zero to a constant concentration for 3s (grey bar) and then set back to 0. OR activation commences immediately upon odour onset and then leads to spiking in ORNs, followed with barely noticeable delay by PN and LN spikes. One can make out a hint of the spike rate adaptation in the example ORN and LN but this becomes more pronounced for higher spike rates. PNs do not have spike rate adaptation in this model.

**Figure 4.**
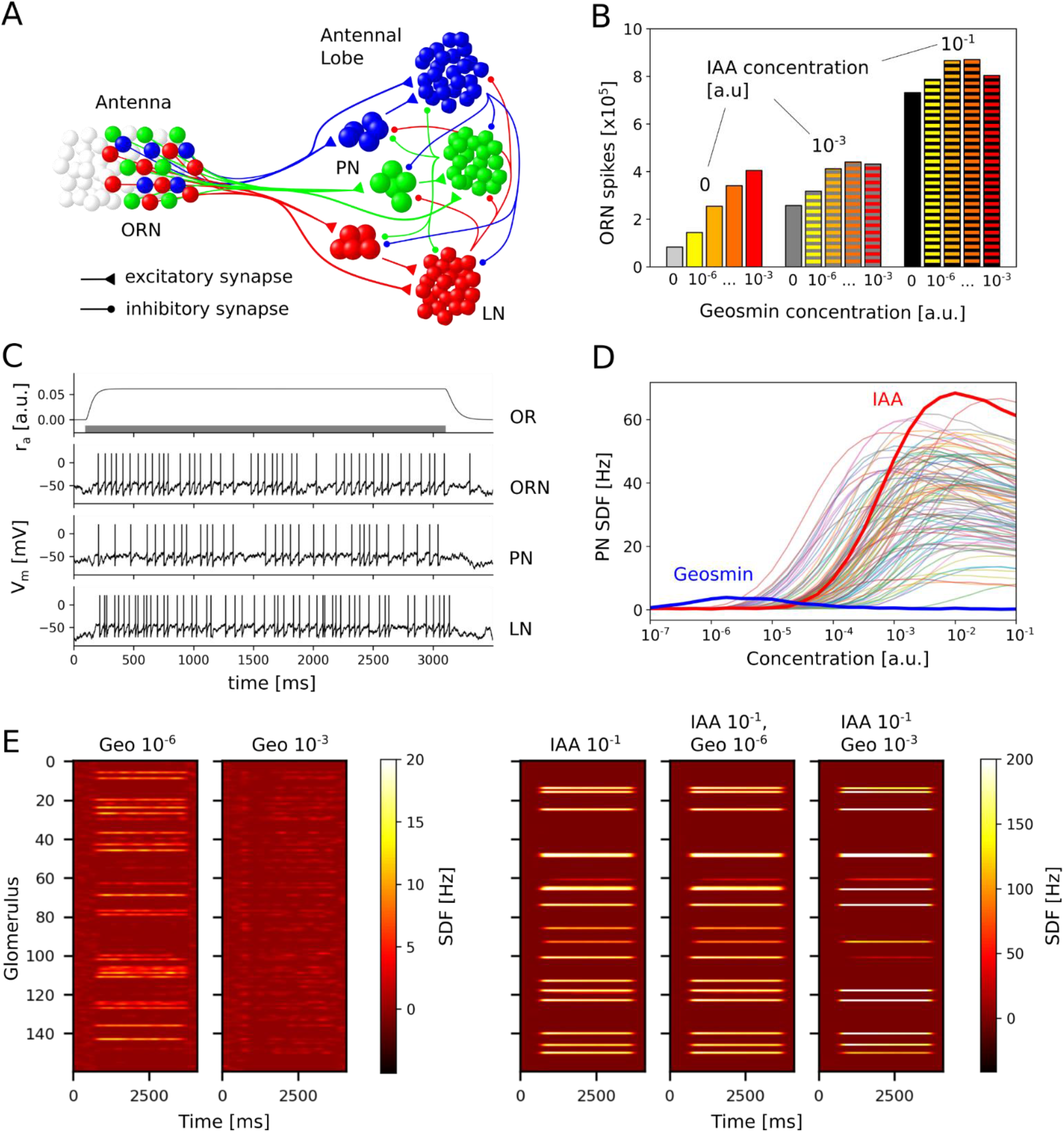
Computational Model. **A.** Circuit diagram of the model. ORNs excite PNs and LNs in the AL according to their receptor type (colours). LNs also receive excitation from the PNs within the same glomerulus. LNs inhibit PNs and LNs in all other glomeruli but not in their own. **B.** Response of the ORNs to different mixtures of the “IAA” and “Geosmin” odours. **C.** Example data for a typical odour response. The bar in the top panel indicates odour exposure. For ORNs, PNs and LNs, the trace of one arbitrary example neuron connected to the most strongly responding glomerulus is shown. Spikes were added to the LIF membrane potential traces as vertical lines for clarity. **D.** Time-averaged responses of the PNs in the strongest responding glomerulus in response to 100 different simulated odours presented for 3s each and at different concentrations. **E.** Heat maps illustrating the mean PN activity across each of the glomeruli in response to “Geosmin”, “IAA”, and mixtures of the two odours.

By inspecting the EAG recordings and PN imaging results we hypothesized that the unusual declining responses for higher concentrations of Geosmin must be due to the local inhibition mediated by LNs in the AL. Furthermore, non-monotonic behaviour has not been widely reported so we reasoned that Geosmin must have a particular property that makes it susceptible to excess inhibition. We explored the properties of odour responses, including the sensitivity (*η* in the OR model), breadth of the response across receptor types (*σ* in the OR model) and the activation (*k_2_* in the OR model). We found that the breadth of the response was the decisive factor leading to non-monotonic response-concentration relationships in the PN. **Figure 4D** illustrates this result. We generated 98 odours with a random distribution of response properties and two with more specific ones, one odour that had a very broad response profile, which we identify with Geosmin, and one odour that had an average width profile but very high activation that we identify with IAA. As can be seen in **Figure 4D**, “Geosmin” shows a non-monotonic response as a function of concentration while “IAA” is essentially monotonic. The random sample of other odours differ in their behaviour but have in the majority typical, increasing sigmoid response curves. We found that the “monotonicity” or lack thereof of odour responses strongly correlates with the width of the odour profile (supplemental **Figures S2–S5**). The intuition behind the non-monotonic behaviour for odours with unusually broad responses is that with increasing concentration, more OR types are activated, increasing the number of involved glomeruli and hence the global inhibition in addition to the increase of inhibition due to increased activation of OR types which are already active. At the same time, the excitation to each glomerulus increases only according to the sigmoid response curve of the corresponding OR type. When the response profile is broad, the combined increases of inhibition can outweigh the increase in excitation and the overall PN response decreases. For narrower response profiles, the excess inhibition from newly recruited OR types is less and the PN response continues to increase.

We then asked whether we can also reproduce the observed interactions of geosmin and IAA seen in the EAG data (**Figure 2**). Here, we interpreted the total number of ORN spikes across all OR types as a sensible proxy of an EAG measurement and reasoned that interactions are likely to be due to syntopic mixture effects at the receptors. With this in mind, Geosmin would lead to inhibition of IAA responses on the antenna if its activation was lower than IAA’s. We, therefore, set the activation rate *k*_2_ of “Geosmin” lower than of “IAA” and generated ORN spike counts for mixtures of “Geosmin” and “IAA”. Through manual exploration, we found that for a roughly 1:3 relationship 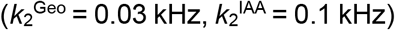 the responses exhibited suppression effects that showed commonalities with the experimental observations (**Figures 2B-D, 4B**). The exact quantitative relationship at different concentration ratios depends besides on the *k*_2_ ratio also on the overlap of “Geosmin” and “IAA” OR activation profiles. We tested this by generating “Geosmin” and “IAA” as Gaussian response profiles with a specific distance between their maxima. For small distances (large overlap), the syntopic suppression of IAA responses dominates, and for large distances (small overlap) the responses are more additive. While the model does reproduce suppression of IAA responses by geosmin in principle, we were not able to fully reproduce the more or less constant responses seen in **Figure 2D** with this scheme and without losing the match of model behaviour and experiments on the level of the AL (see above). It is, however, likely that with other ways of generating overlapping response profiles or, indeed, other than Gaussian response profiles, the experimental observations could be replicated more closely.

Finally, we asked whether the “Geosmin” and “IAA” odours generated to reproduce the non-monotonic behaviour in PNs and the sub-additive response properties in the EAG would produce PN responses that are similar to the calcium imaging data of PN activity in the AL. **Figure 4E** shows the simulated response patterns of the PNs in response to Geosmin at 10^−6^, 10^−3^ (compare **Figure 3A**), IAA at 10^−1^, and the mixtures of IAA at 10^−1^ and Geosmin at 10^−6^ and 10^−3^ (compare **Figure 3B**). Overall, the responses look similar to the experimental data with a moderate response to “Geosmin” 10^−6^, which essentially disappears at 10^−3^. In the mixtures, adding “Geosmin” 10^−6^ to IAA 10^−1^ has almost no visible effects while adding “Geosmin” 10^−3^ appears to sharpen the response profile somewhat, where glomeruli strongly activated by “IAA” and “geosmin” get even more activated, and weakly activated glomeruli are slightly depressed, presumably due to added global inhibition by the LNs. By visual inspection, similar effects appear to be present in the experimental data (**Figure 3B**) but we did not identify statistically significant effects.

## Discussion

### Behavioural Effects

Our initial hypothesis, that geosmin would increase stinging responses - similar to what is observed in environments with high humidity (Brandeburgo, M. M Goncalves and Kerr, 1982; Southwick and Moritz, 1987), the condition in which geosmin is released by soil bacteria (Liato and Aïder, 2017) - was not confirmed: geosmin by itself did not affect the stinging likelihood of honey bees in our assay. Our results even demonstrate the opposite effect, whereby the presence of geosmin inhibits the defensive response usually elicited by the major component of the honey bee alarm pheromone (**Figure 1**). Intriguingly, we tested the bees with 2 concentrations of geosmin and only found this inhibiting effect when using the lowest one. Concentration-dependent effects of odours on behaviour have been reported before, for example weaver ants exhibit a series of behaviours as they approach (and thus perceive an increase in concentration) a source releasing 1-hexanol, a component of their own alarm pheromone (Bradshaw et al., 1979). Recently, non-monotonic behavioural responses to IAA itself have been reported in honey bees (López-Incera et al., 2021). More precisely, the stinging frequency of individual bees tested in the same behavioural assay increased with IAA concentration up to 2.5∙10^−1^, but then dropped back when higher concentrations were used. Added to the strong overlap observed between the glomeruli activated by geosmin and IAA, one hypothesis could be that additive activation by geosmin would shift the behavioural response towards the declining range of this IAA dose-response curve. While this hypothesis needs to be further tested, we find it unlikely since the decrease in stinging behaviour only appears at very high IAA concentrations, presumably requiring much stronger activations than the ones recorded here.

The modulation of the defensive behaviour observed here is similar to that of appetitive floral odours (Nouvian et al., 2015), which may suggest that geosmin has a positive valence for honey bees. To test this hypothesis, one could measure the frequency of spontaneous extension of the proboscis in response to this compound, as previously done (Nouvian et al., 2015). *Streptomyces* bacteria have been found in flowers and can protect honeybees against pathogens (Kim et al., 2019). While the vast majority of *Streptomyces* species can produce geosmin (Yamada et al., 2015), actual detection of geosmin within the floral bouquet has however only been reported in a few cacti species (Kaiser, 2004). Whether this compound could attract bees and thus participate in mediating mutual interactions thus remains to be verified. Geosmin is an ecologically relevant compound eliciting either attraction or repulsion in a number of arthropod species including flies (Becher et al., 2010; Stensmyr et al., 2012), mosquitoes (Rodríguez-Hernández et al., 2019) and springtails (Becher et al., 2020). Although its ecological function remains mysterious in the case of honey bees, our data suggest that it may be equally relevant for this important pollinator, and worth investigating further.

### ORN responses

Electroantennography results suggest that the non-monotonic concentration dependence of the behaviour does not stem from an anomalous sensitivity of receptors to geosmin, as the EAG amplitude shows a typical increasing dose-response curve for stimuli with the pure odour (**Figure 2B**). However, since a behavioural effect was observed only to mixtures between geosmin and IAA, most interesting are potential interactions between these two signals. Indeed, compared to an expectable linear sum of the pure odour response amplitudes (Akers and Getz, 1993), the response amplitude to the binary mixture is not showing this clear increase with concentration. In a mixture with low IAA concentration (**Figure 2C**), the addition of geosmin 10^−6^ does not increase the signal as could be expected (Akers and Getz, 1993). For the high IAA concentration (**Figure 2D**), the effect is even more pronounced: the addition of geosmin 10^−5^ causes a significant decrease in the signal, and the expected increase is not observable until a geosmin concentration of 10^−3^. Dose-response curves in which a higher concentration does not necessarily produce an increase in the response amplitude, have been observed previously in EAG measurements in honey bees, and these were also responses to odour mixtures (Iwama et al., 1995).

An interaction of geosmin and IAA is in line with the behavioural signal, where the effect of geosmin on IAA-induced stinging was significant for geosmin 10^−6^ but vanished for geosmin 10^−3^.

The underlying mechanism for this interaction could be a masking of IAA by geosmin at the level of the receptors, an effect that has been reported (Kurahashi et al., 1994). Interestingly, in this work it was IAA that had been identified as a strong masking agent but was also shown to be maskable by some odours (Kurahashi et al., 1994). One requirement for masking is a sensitivity of receptors to both odours. This was one of the questions to be clarified by resolving the integrated neuronal response that the EAG signal provides by means of calcium imaging of the odour response maps at the level of the antennal lobe.

### PN responses

This hypothesis was indeed confirmed by the response maps of the glomerular projection neurons. Although only a subset of 19 out of 160 glomeruli (**Figure S1**) could be consistently identified in various subjects, all the glomeruli that were activated by IAA were found to be also activated by geosmin 10^−6^ (**Figure 3**).

A further surprising finding of the calcium imaging was a non-monotonic dependence of the projection neuron responses to the geosmin concentration (**Figure 3A, B**). Using the same concentrations as in the behavioural studies, the broad and strong responses to geosmin 10^−6^ almost completely disappeared for 10^−3^. This again is in line with the behavioural data. However, such a non-monotonic concentration dependence seems to be very rare. At the level of the antennal lobe, it was not reported in bees before. A work in moths shows such responses to pheromone components from electrophysiological recordings in the AL, the type of neurons however remains undetermined (Varela et al., 2011). At the level of the honey bee mushroom body input, PN boutons showed highly varying concentration dependencies (Yamagata et al., 2009), monotonic increases and decreases as well as non-monotonic changes. The authors suggest inhibition at the level of the boutons. Another work in moths found only monotonic increases at the level of the AL in the PN dendrites, but at the level of the PN somata also non-monotonic responses (Fujiwara et al., 2014). The suggested underlying mechanism was postsynaptic inhibition. In addition to those two positions at which inhibition could create such a change in concentration dependence, our data now suggest that also inhibition within the antennal lobe might have an effect.

Imaging the response to mixtures of IAA and geosmin confirmed the non-monotonic dependence on the geosmin concentration also under this condition. However, an interaction of IAA and geosmin was not observed in the imaged subset of glomeruli, the mixtures elicited a response that appears to be the exact sum of the pure odour responses of both compounds (**Figure 3B**). This suggests two possible scenarios. Either the strength of this interaction is varying across glomeruli and is not visible in the subset that we were able to image, or the observed behaviour is not induced by an interaction of both odours at the level of their identification in the antennal lobe, but when the valence of these odours is extracted in the higher brain centres like the mushroom body and the lateral horn (Li and Liberles, 2015).

A final important result of the calcium imaging is the observation of a broad response map that geosmin elicits, which proves combinatorial coding of this odour in contrast to the results in fruit flies (Stensmyr et al., 2012) and mosquitoes (Melo et al., 2020) where the response was limited to a single glomerulus, which corresponds to the complementary coding mechanism of a labelled line (Hildebrand and Shepherd, 1997). Studies of the evolution of olfactory receptors indeed suggest that a variation in coding modes is likely between species (Andersson et al., 2015). The width of the spectrum of receptors that geosmin activated might be even slightly broader than those of the floral odours that were imaged in comparison (**Figure 3A**), which was one indication of why the anomalous concentration dependence was found.

### Neuronal network model

The puzzling results of a non-monotonic concentration dependence of the geosmin response in the antennal lobe output neurons, as well as the interaction between geosmin and IAA at the level of the olfactory receptor neurons, were discussed with the help of a computational model of the AL and we have demonstrated that such a response pattern can arise as a result of a specific balance of excitatory-inhibitory coupling in the AL. Furthermore, the unusual concentration tuning of this response suggests that geosmin processing may be different from that of other odours and the model predicts that, in contrast to other insects, geosmin responses in bees are indeed broader rather than narrower than other odours. Taken together, our data and model provide a first evidence that geosmin is a differentially processed and ecologically relevant odorant for honey bees.

### Outlook and Conclusions

These results constitute another step towards understanding olfactory modulation of defensive behaviour in honey bees, both at the behavioural and neuronal levels. Besides a fundamental understanding of the neuronal circuits leading to stinging, this may also be of practical use for beekeeping.

Recent studies proposed the use of geosmin as a natural repellent against pest insects such as *D. suzukii* (Wallingford et al., 2017). Further investigation on the consequences of geosmin on non-target insects such as honey bees should be performed before any commercialization of geosmin for this purpose.

Studies should also be extended to guard bees, which could show an even stronger contrast in the interaction between IAA and geosmin to further test the hypotheses made here. Context is key for pheromone response, hence future experiments at the hive level will be required to fully understand the effect of geosmin.

Following the elicited neuronal activity into the higher-order olfactory processing centres such as the lateral horn and mushroom bodies would also be a valuable future direction in order to study the valence of geosmin, IAA, and its mixtures at different concentrations. This might be complemented with additional modelling to better underpin our understanding of underlying circuits and processes and generate testable hypotheses to challenge that understanding.

Finally, the original hypothesis of geosmin as a weather indicator should be followed up. This could help to anticipate problems in the adaptation of bees to climate change and thus prevent ecological and economic damage.

## Material and Methods

### Behavioural experiments

#### Honey bees and preparation procedure

Honey bee foragers were collected from various colonies of *Apis mellifera ligustica*, located in Rovereto, Italy from September 2019 to November 2019 and July 2020 to August 2020. The colonies were freely foraging and underwent routine beekeeping inspections during the entire period of the experiments. An equal number of bees from different colonies were included in the behavioural experiments. The bees were caught on sunny and cloudy days (but not on rainy days) in two rounds (around 10:30 AM or 14:00 PM).

Foragers were collected using a plastic container as they exited the hives, and they were brought back inside the lab and placed in an icebox. When the bees were motionless, they were placed in pairs into 50 ml centrifuge tubes modified into syringes. Two droplets of sucrose solution (50% sucrose water, vol/vol) were placed into the tube after the bees recovered completely. All honey bees were allowed to recover for at least 15 min (up to ~1h for the last bees) before being tested in the set-up investigating stinging behaviour. If one or both bees showed signs of poor recovery when put in the setup (difficulty to hold upside down, disorientation and/or lethargic walk) the whole trial was excluded from further analysis. All the materials used to contain the bees were washed and cleaned with 80% ethanol, before the next use.

In total, 288 bees participated in the behavioural experiments, equally distributed between the 6 odour conditions (hence a sample size of 48 bees per group). This sample size was chosen based on previous studies (Nouvian et al., 2015, 2018).

#### Odour stimuli

All odours were obtained from Sigma-Aldrich (98–99.9% purity) and were stored at 4°C. They were diluted in mineral oil at the start of the experimental period and kept for the whole length of the behavioural experiment. When not in use these odours were sealed and stored at room temperature. The main component of the sting alarm pheromone, isoamyl acetate (IAA) was diluted to 10^−1^ (vol/vol) as in previous studies (Nouvian et al., 2015, 2018). The concentrations of 10^−3^ and 10^−6^ were chosen for geosmin as these were shown to elicit behavioural responses in fruit flies (Stensmyr et al., 2012) and mosquitoes (Melo et al., 2020).

The odours were delivered at room temperature (24°C) by placing a filter paper soaked with 10 *μ*l of odorant solution into an airflow that was injected into the testing arena. The flow remained on during the whole duration of the trial (3 min). For testing interactions between 2 odours, 2 filter papers each carrying one of the odours were placed into the airflow.

#### Stinging assay

The bees’ stinging responsiveness was tested using the assay described in detail in Nouvian et al., 2015. Briefly, dyads of honey bees were introduced into a cylindrical testing arena in which they confronted a black rotating dummy, prolonged by a black feather (**Figure 1A, Video 1**). The primary function of the black feather is to disturb the bees without causing any pain, by brushing the sides of the arena. Note that the bees can easily avoid both dummy and feather. A trial lasted 3 min and was scored as “stinging” if at least one of the bees decided to sting the dummy during this time. This behaviour was defined as the bee holding onto the dummy or the feather for at least 3 seconds, with the tip of the abdomen pressed against it in the characteristic stinging posture. All the behavioural trials were recorded with a web camera (Microsoft Life cam) placed above the arena.

Two identical arenas and 2 dummies were used. Their use was balanced across the different odour conditions, to ensure that they did not contribute to potential differences in behaviour. Before each trial, the arena and the dummies with their feathers were cleaned using an 80% ethanol solution.

#### Data analysis

A generalized linear model (glm) was used to analyse the percentage of stinging trials. The odour group was set as a fixed factor while the hive and dummy were defined as random factors. A pairwise comparison (glht package in R) was done followed by Benjamini and Hochberg false discovery rate (FDR) correction for controlling Type I errors.

### Electroantennography (EAG)

#### Preparation procedure

The EAG technique was adapted from Anfora et al., 2010, and the recordings were performed using a standard EAG apparatus (Syntech, Hilversum). Honey bee foragers were collected and handled in the same way as for the behavioural assay. After chilling them on ice, one antenna of each animal was cut at the level of the scape. A total of *n=24* antennas from both left and right sides were used for this experiment (14 left and 10 right antennas) to avoid lateralization effects (Haase et al., 2011). The base of each antenna was then inserted into the glass reference electrode filled with Kaissling saline solution (Kaissling and Thorson, 1980). The recording electrode was brought into contact with the last segment of the flagellum from which the distal tip had been cut.

#### Odour stimuli

A custom-made olfactometer was used to deliver odours to the bee antennas. The odour stimuli originate from glass vials containing 1mL of odours dissolved in mineral oil. The olfactometer was operated using LabView and the single channels were switched by solenoid valves (LHDA0531115, The Lee Company) controlled by a PCIe-6321 multifunction board (National Instruments). The airflow during the recording is maintained constant during all phases of the experiment (For details see Paoli et al., 2017).

Seven mineral oil solutions of either Geosmin (Geo) at concentrations 10^−6^, 10^−5^, 10^−4^, 10^−3^ vol/vol, Isoamyl Acetate (IAA) at concentrations 10^−3^, 10^−1^ vol/vol, or pure mineral oil (control) were prepared. Olfactometer glass vials were filled with these solutions. The stimulation protocol for each bee consisted of presenting each odour and combination of odours in ascending order of concentrations, giving rise to 15 stimuli, see **Figure 2B-D**: Control, Geo 10^−6^, Geo 10^−5^, Geo 10^−4^, Geo 10^−3^, IAA 10^−3^, IAA 10^−3^+Geo 10^−6^, IAA 10^−3^+Geo 10^−5^, IAA 10^−3^+Geo 10^−4^, IAA 10^−3^+Geo 10^−3^, IAA 10^−1^, IAA 10^−1^+Geo 10^−6^, IAA 10^−1^+Geo 10^−5^, IAA 10^−1^+Geo 10^−1^, IAA 10^−1^+Geo 10^−3^. The stimulus duration was 2 s and the inter-stimulus interval was 20 s. In addition, each cycle was repeated 10 times and therefore a total of 150 stimuli were presented to each bee antenna, this resulted in recordings of 55 min total duration.

#### Data analysis

The response amplitude was calculated by subtracting the voltage averaged during 1 s before each stimulus from the voltage averaged during 1 s after the beginning of the stimulus. These values were thereafter averaged over the ten repetitions of each stimulus. Responses were analysed via repeated-measures ANOVA followed by Bonferroni post-hoc tests.

### In vivo calcium imaging

#### Preparation procedure

Honey bees were prepared for the *in vivo* calcium imaging experiment according to a well-established protocol (Paoli et al., 2017). The foragers used were collected and immobilized in a fridge at 4°C for 5-6 min. The immobilized bees were then fixed onto a custom-made imaging stage, using soft dental wax (Deiberit 502, Siladent). A small rectangular window was cut into the head cuticula of the fixed bee. The glands and trachea were moved aside and fura2 - dextran, a calcium-sensitive fluorescent dye (Thermo-Fischer Scientific) dissolved in distilled water was injected into the antenna-cerebralis tracts right below the α-lobe using a microtip (Paoli et al., 2016). After the injection, the cuticula was fixed in its original state using n-eicosane. The bees were stored in a dark, cool, and humid place for about 20 h to ensure that the calcium dye has diffused into the AL.

Just before the imaging session, the cuticular window, trachea, and glands were completely removed. A silicone adhesive (Kwik-Sil) was used to cover the open window on top of the bee’s head, and it was left to dry for a few minutes. The fluorescent signal in the antennal lobe was then imaged under a two-photon microscope.

#### Two-photon microscopy

The two-photon microscope (Ultima IV, Bruker) is based on an ultra-short pulsed laser (Mai Tai, Deep See HP, Spectra-Physics). The laser was tuned to 780 nm for fura-2 excitation. All images were acquired with a water-immersion objective (10×, NA 0.3, Olympus). The fluorescence was collected in epi-configuration, selected by a dichroic mirror, and filtered with a band-pass filter centred at 525 nm and with a 70 nm bandwidth (Chroma Technology Corp). Finally, it was detected by a photomultiplier tube (Hamamatsu Photonics). Laser powers of about 10 mW were used in order to balance signal to noise ratio (SNR) against photo-damage effects that reduced the bee life span.

The field of view of 280 × 280 μm^2^ was resolved by 128 × 128 pixels. The fluorescence intensity was recorded with a depth of 13 bits. The image acquisition at a frame rate of 10.1 Hz was synchronized to the stimulus protocol.

In addition to the functional images, a *z*-stack of the antennal lobe was acquired with a spatial resolution of 512 × 512 pixels and a z-layer distance of 2 μm to perform the morphological identification of glomeruli.

#### Odour stimuli

The olfactometer used to deliver odours under the two-photon microscope was the same used in the EAG experiment. During an imaging session, the odorants of interest (Geosmin 10^−6^, Geosmin 10^−3^, and 10^−1^ IAA) were presented to the bee in a sequence either as a single odour and as mixtures, and the sequence was repeated 10 times. Each stimulus pulse lasted 3 s with a 12 s inter-stimulus interval. For comparison of response strength and width also 3 floral odours were tested with the same sequences (1-nonanol 5∙10^−3^, benzaldehyde 5∙10^−3^, and 3-hexanol 5∙10^−3^)

#### Data post-processing and analysis

A total of 14 bees were recorded and analysed. Data analysis was automatically executed after the experiments employing custom scripts in MATLAB (R2019b, MathWorks). From the structural data and with the help of the antennal lobe atlas (Deisig et al., 2006), individual glomeruli were identified and assigned to regions of interest in the functional data. Averaging the fluorescence signal over each ROI provided the raw data for the individual glomerular responses. These time series were then separated into periods of 3 s pre-stimulus, 3 s during stimulus and 3 s post-stimulus for each trial. For each frame we computed the relative fluorescence change Δ*F*/*F* = −[*F*(*t*) - *F_b_*]/*F_b_* where *F_b_* is the average fluorescence signal in the pre-stimulus period. This signal is proportional to the relative change in calcium concentration and therefore the neuronal firing rate (Moreaux and Laurent, 2007). Finally, Δ*F*/*F* was averaged over the 10 trials producing glomerular response curves in each bee to all odour stimuli in all identified glomeruli.

#### Computational Model

The neurons in the model are described by an adaptive leaky integrate-and-fire neuron with membrane potential equation

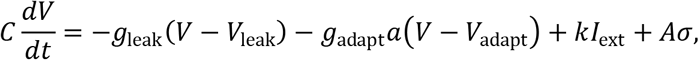

where 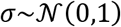 is normally distributed white noise and the factor *k* is used to rescale input currents in the ORNs. For olfactory receptor neurons (ORNs) the input current is the current passing through olfactory receptors (ORs) and *k* = 10, while for local (LN) and projection neurons (PN), it is the synaptic currents from incoming synapses with *k* = 1. All neurons have *C* = 1 nF, *V*_leak_ = −60 mV, *g*_leak_ = 10 nS, and *A* = 1.4 nA. Whenever the membrane potential *V* crosses the firing threshold *V*_*th*_ = −40 mV, a spike is emitted and the membrane potential is reset to *V*_reset_ = −70 mV. PNs do not have an adaptation current, but ORNs and LNs have spike rate adaptation with *g*_adapt_ = 1.5 nS for ORNs, and *g*_adapt_ = 0.5 nS for LNs. The adaptation variable was governed by

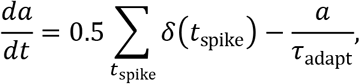

where *δ* is the Dirac delta distribution and *τ*_adapt_ = 1s for both ORNs and LNs.

Synapses are described with an instantaneous rise of synaptic activation *s* upon arrival of a spike and subsequent exponential decay,

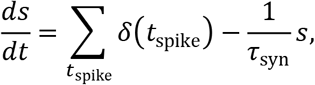

and a conductance-based input current into the post-synaptic neuron,

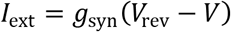

with reversal potential *V*_rev_ = 0mV for excitatory (ORN to PN, ORN to LN, PN to LN) and *V*_rev_ = −80mV for inhibitory (LN to PN and LN to LN) synapses.

Olfactory receptors and the process of transduction are described by a standard two-stage binding and activation rate model,

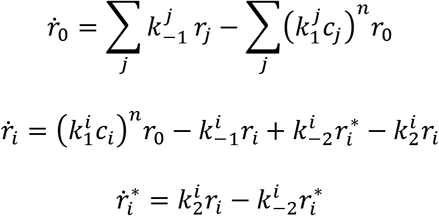

Where *r*_0_ is the fraction of unbound receptors, *r*_*i*_ the fractions of receptors bound to odours *i* = 1, …, *N*, and 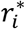 are the fractions of receptors bound to and activated by odours *i*. The constants 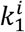, and 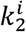 respectively describe the rate of binding to and being activated by odour *i*, while 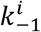, and 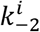 describe the of unbinding and inactivation. All *k* constants can be specific to the odours and receptor types. In this work, we chose individual constants 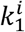 for each odour-receptor type pair and odour specific (but equal for all receptor types) 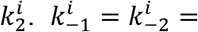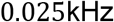 were identical for all odours and receptor types.

The individual binding constants for each odorant across receptors were chosen as Gaussian profiles,

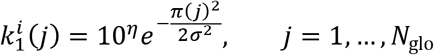

where *π*(∙) is a (randomly chosen) permutation of 1, …, *N*_glo_, and *η* is a 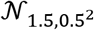 distributed random variable, truncated to values within [0, 4]. The standard deviation *σ* of the odour profiles was sampled from 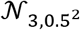, truncated to values greater or equal 1.5, all in units of kHz. The activation constant 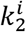 for each odour was sampled from 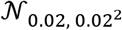, truncated to values within [0.0028, 0.2], in units of kHz.

For the simulations in this work, we generated 98 odours according to these rules and two additional odours, which we identify with IAA and Geosmin. For “IAA”, we used *η* = 0.8, *σ* = 3, and *k*_2_ = 0.1. For “Geosmin”, *η* = 4.4, *σ* = 10, and *k*_2_ = 0.003, which is the combination of a broad binding profile, small activation and high sensitivity. We generated the “IAA” and “Geosmin” odours with a fixed distance of 30 OR types from peak to peak before scrambling both with the same permutation. This ensures that the two odours have a specific amount of overlap with implications in particular for the interactions at the OR on the antenna, as evidenced through the ORN spike counts.

The model network is illustrated in **Figure 4A**. We simulated 160 receptor types and 60 ORNs for each type. Each of the 5 PNs and 25 LNs in a glomerulus are connected to 12 randomly chosen ORNs of their receptor type. PNs excite all LNs in their glomerulus and LNs inhibit all PNs and LNs in all glomeruli except their own. The synapse parameters for these connections are summarised in **Table 1**. All differential equations were integrated with a linear Euler algorithm and global 0.2ms timestep.

**Table 1:** Synapse parameters

|  | ORN→PN | ORN→LN | PN→LN | LN→PN | LN→LN |
| --- | --- | --- | --- | --- | --- |
| $g_{\text{syn}}$ (nS) | 8 | 8 | 1 | 0.055 | 0.02 |
| $\tau_{\text{syn}}$ (ms) | 10 | 10 | 10 | 20 | 20 |
| $V_{\text{rev}}$ (mV) | 0 | 0 | 0 | -80 | -80 |

The models were implemented using the PyGeNN interface (Knight et al., 2021) for GeNN 4.5.0 (Yavuz et al., 2016; Knight and Nowotny, 2018). GeNN is available at ext-link https://github.com/genn-team/genn and the source code for the modelling work in this paper is available at https://github.com/tnowotny/bee_al_2021, including Jupyter notebooks for analysis and plotting. Simulations were run on a Linux workstation, running Ubuntu 18.04.5 LTS.

## Acknowledgements

F.S. was supported by MIUR, project PRIN 2017MKNP2F. E.T. was supported by the strategic project BRANDY of the University of Trento. A.C. was funded by the Autonomous Province of Bolzano (Project B26J16000310003). M.N. was supported by funding from the Zukunftskolleg (University of Konstanz) and the Erasmus + Staff mobility program. T.N. was supported by the EPSRC (EP/P006094/1, EP/S030964/1), a Leverhulme Trust Research Project Grant and the European Union’s Horizon 2020 research and innovation program under grant agreement No 945539 (HBP).

The authors declare no competing financial interests.

---

**Video 1.**
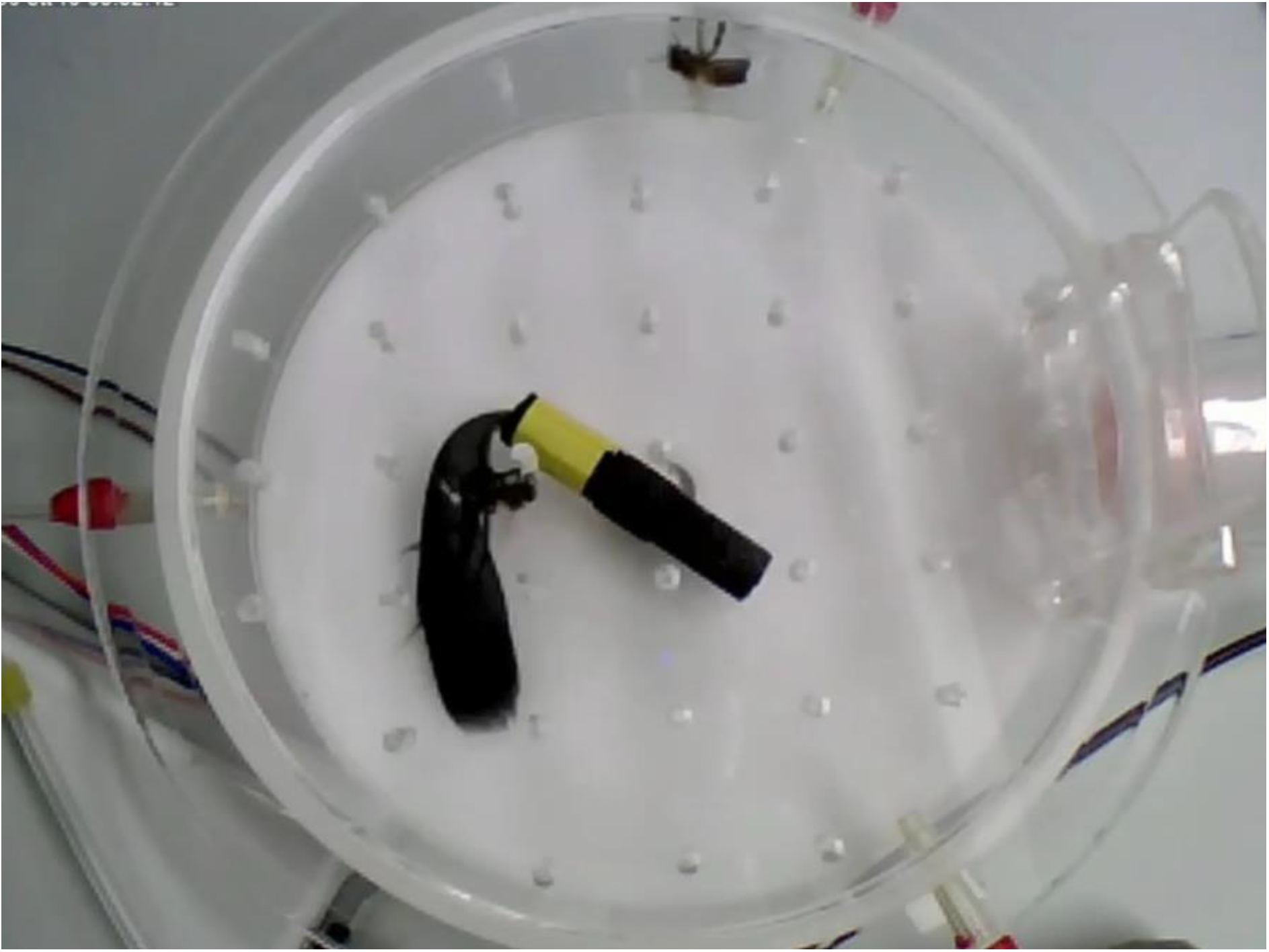
Initial phase of the stinging assay. A Dyad if bees is inserted into the experimental arena, one of them immediately attacks the rotating dummy.

## Supplementary information

**Figure S1.**
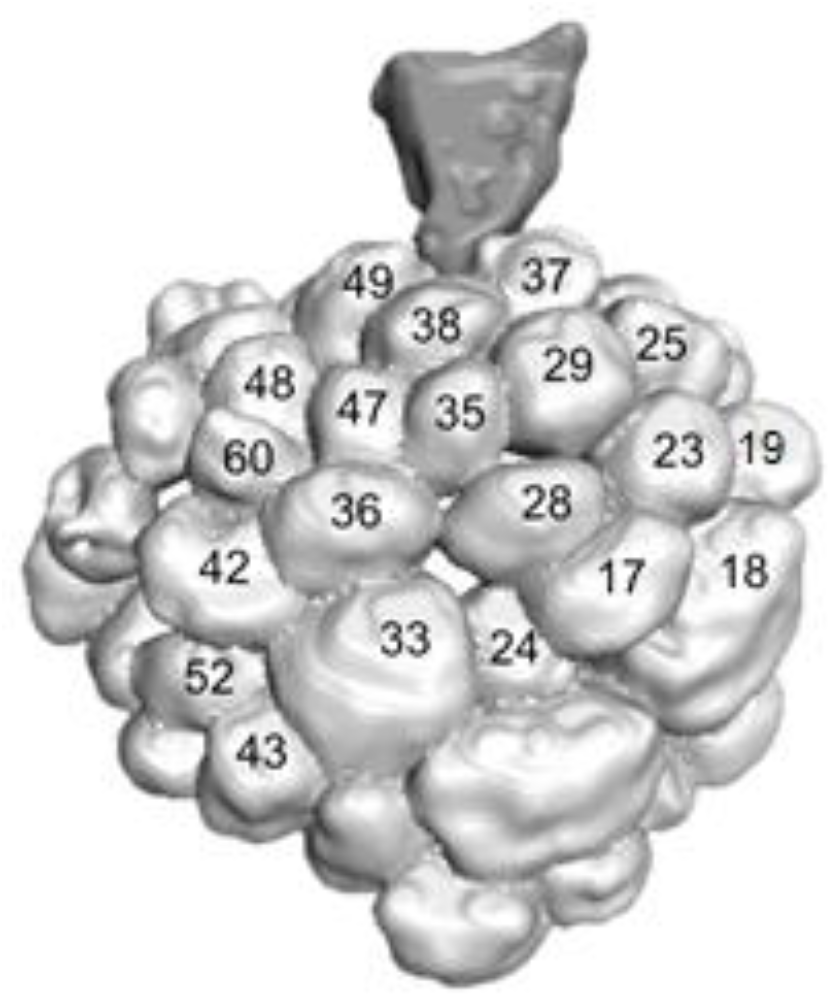
Anatomical map of the antennal lobe. Labels mark the 19 responsive glomeruli. Image adapted from (Paoli et al., 2016).

**Figure S2.**
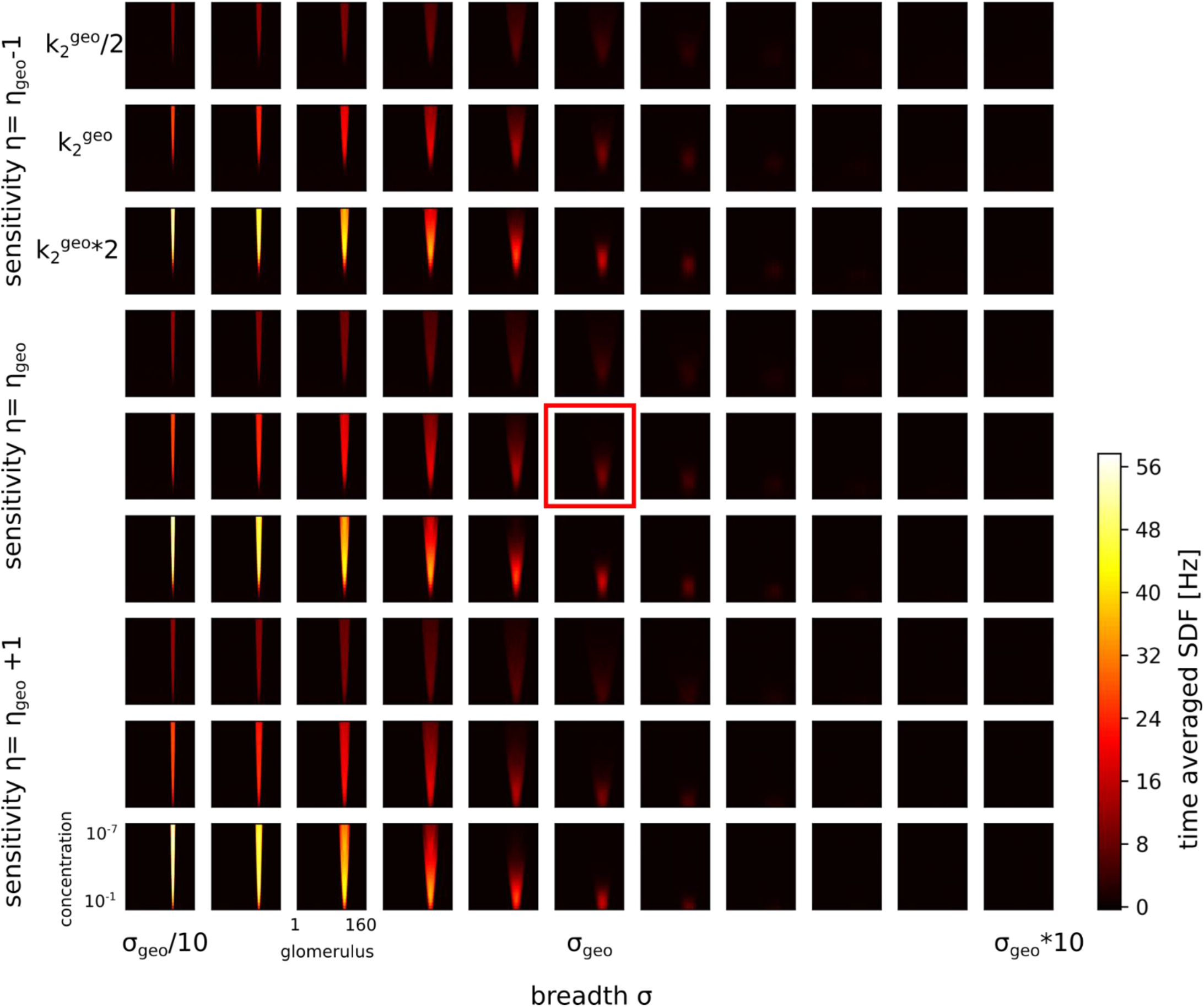
Odour responses in PNs as a function of concentration. For a selection of 3 different properties of the odour profile: (i) the breadth *σ* of the odour profile, the overall sensitivity *η* of the ORs to the odour, and the activation *k_2_* of the ORs. The colourmaps all use a common colour scheme as illustrated by the colour bar. The panel with the red rectangle is the “geosmin” odour. Note how the non-monotonic responses start appearing for increasing breadth (from left to right) and how for very broad profiles the responses disappear.

**Figure S3.**
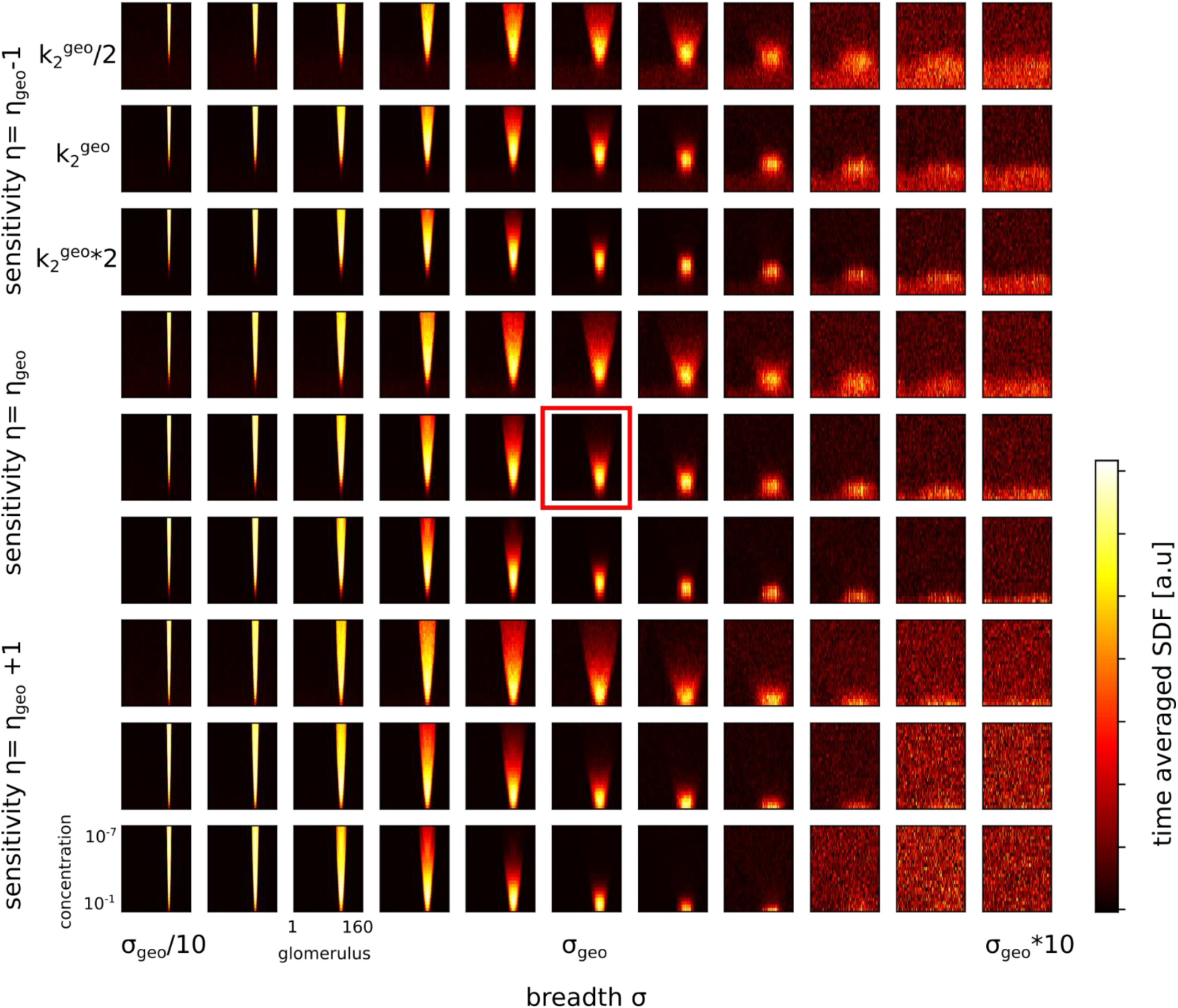
Same as Supplementary **Figure S2** but with individual colour scale for each sub-panel to reveal the details of the spiking activity.

**Figure S4.**
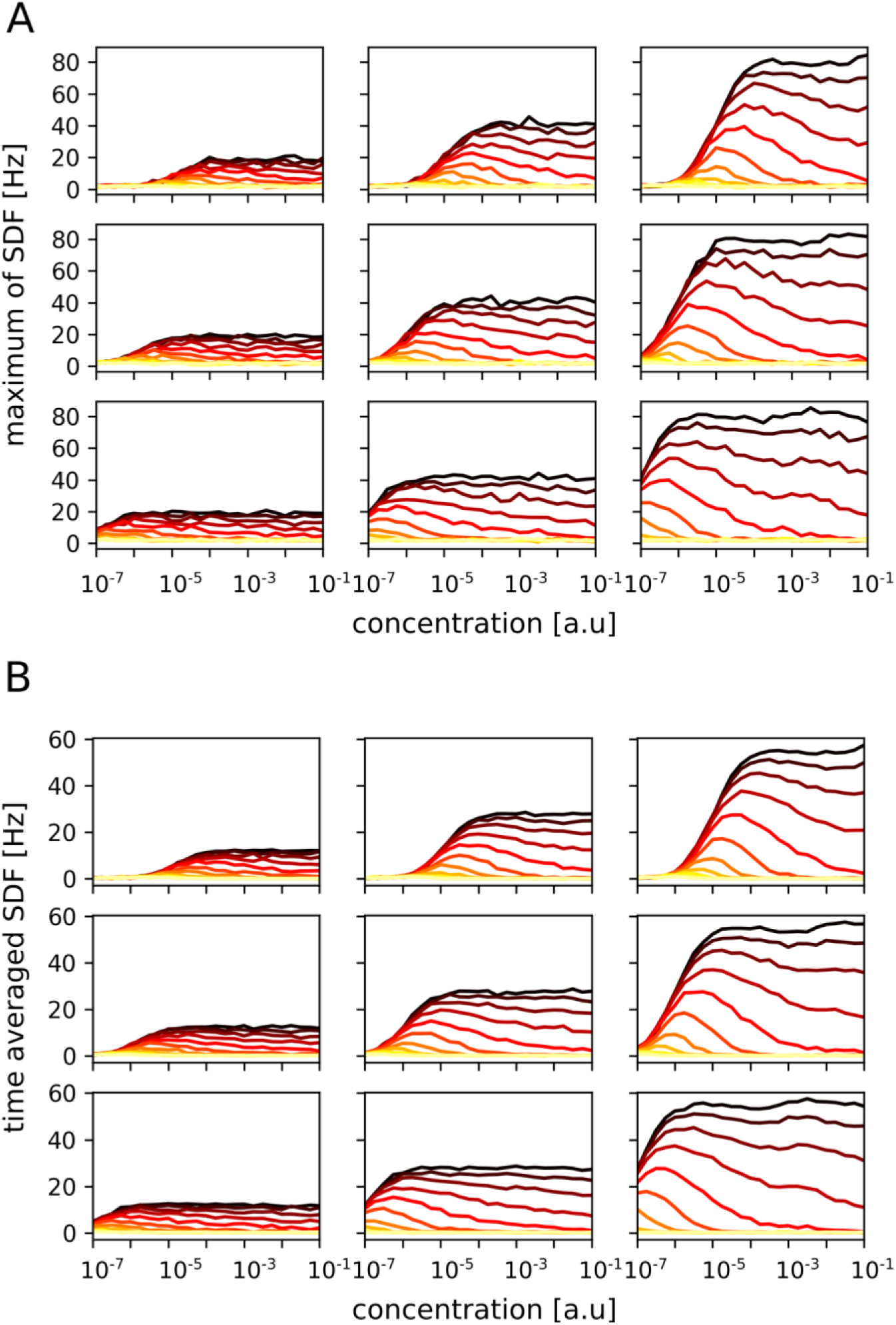
Response of the strongest glomerulus. Dependence of the response of the PNs of the strongest glomerulus as a function of concentration (*x*-axes). **A.** Maximal PN response in the most strongly responding glomerulus. **B.** Time-averaged response of the PNs in the most strongly responding glomerulus. The individual panels correspond to activation *k_2_^geo^*/2, *k_2_^geo^*, *k_2_^geo^*∗2 (left to right) and sensitivity *η_geo_*−1, *η_geo_*, *η_geo_* +1 (top to bottom). The coloured lines are for increasing odour profile breadth *σ*from *σ*_*geo*_/10 (black) to *σ*_*geo*_∗10 (bright red).

**Figure S5.**
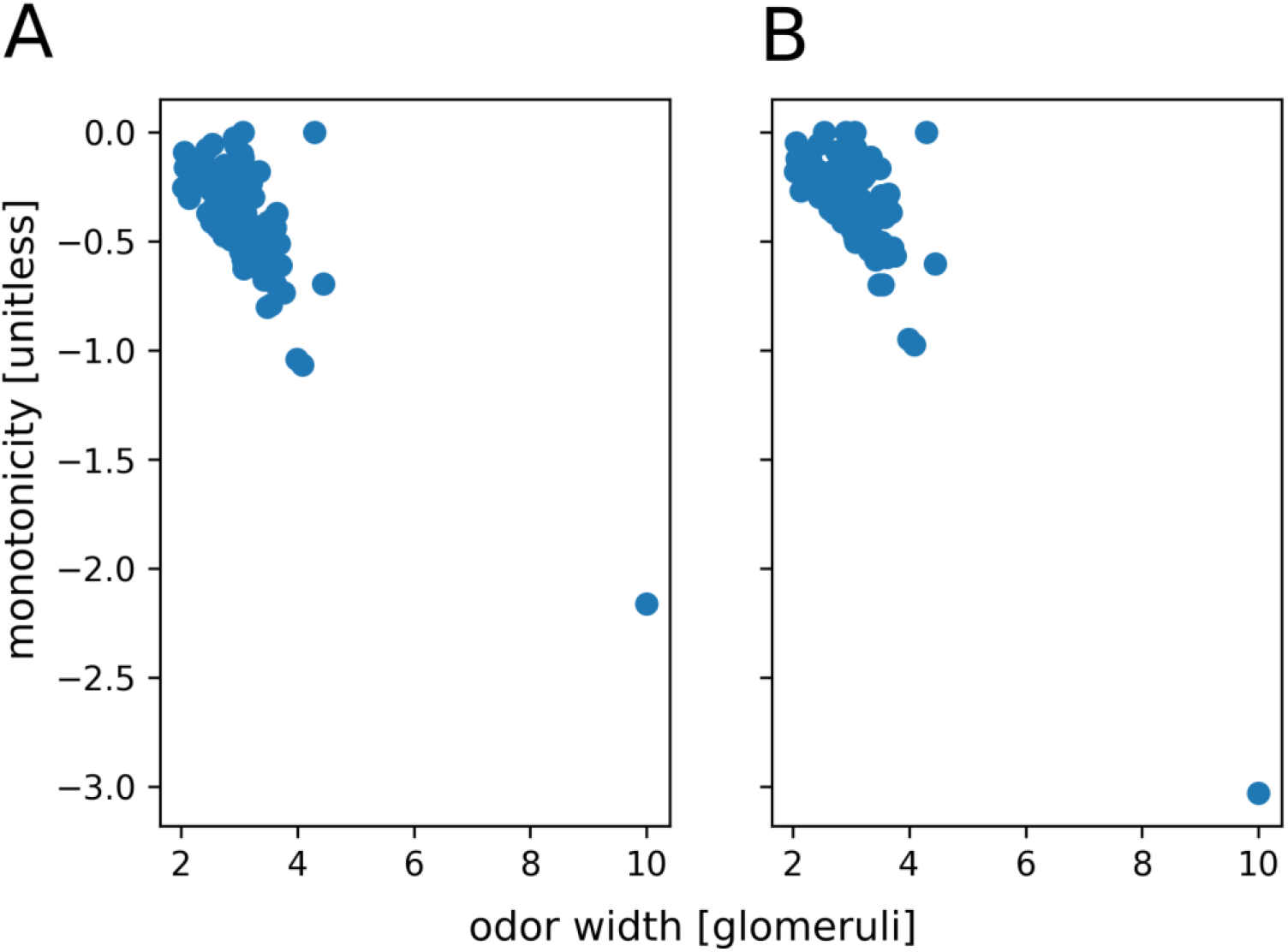
Monotonicity-width relation. Relationship between the width of the odour profile and the observed monotonicity of the response of PNs. **A.** Monotonicity calculated with respect to the maximal (phasic) response and **B.** Monotonicity calculated with respect to the time-averaged response during odour presentation. The monotonicity of the response *x* (*x* = maximal SDF or averaged SDF) in this context was calculated as m = (*x* (10^−1^) - max(*x*))/mean(*x*), where the maximum and mean were taken across all concentrations from 10^−7^ to 10^−1^. Monotonicity takes values <=0, where monotonic odours have *m* = 0. Monotonicity is strongly anti-correlated with odour profile width with *R* = −0.802 for the maximal SDF (A) and *R* = −0.862 for the average SDF (B).

